# Taurine or *N*-acetylcysteine treatments prevent memory impairment and metabolite profile alterations in the hippocampus of high-fat diet-fed female mice

**DOI:** 10.1101/2022.02.02.478774

**Authors:** Alba M. Garcia-Serrano, Joao P.P. Vieira, Veronika Fleischhart, João M.N. Duarte

**Affiliations:** Department of Experimental Medical Science, Faculty of Medicine, Lund University, Lund, Sweden; Wallenberg Centre for Molecular Medicine, Lund University, Lund, Sweden

**Keywords:** brain metabolism, obesity, diabetes, glucose, dementia, neuroprotection

## Abstract

Obesity impacts brain metabolism and function, and constitutes a risk factor for dementia development. Long-term exposure to obesogenic diets leads to taurine accumulation in the hippocampus of rodents. Taurine has putative beneficial effects on cell homeostasis, and is reported to improve cognition. Therefore, hippocampal taurine accumulation in obese and diabetic models might constitute a counteracting response to metabolic stress. This study tested the hypothesis that treatment with taurine or with *N*-acetylcysteine (NAC), which provides cysteine for the synthesis of taurine and the antioxidant glutathione, prevent obesity-associated hippocampal alterations and memory impairment. Female mice were fed either a regular diet or a high-fat diet (HFD). For the same period, some mice had access to 3%(w/v) taurine or 3%(w/v) NAC in the drinking water. After 2 months, magnetic resonance spectroscopy was used to measure metabolite profiles. Then, memory was assessed in novel object and novel location recognition tests. HFD feeding caused memory impairment in both tests, and reduced concentration of lactate, phosphocreatine-to-creatine ratio, and the neuronal marker *N*-acetylaspartate in the hippocampus. Both taurine and NAC prevented HFD-induced memory impairment and *N*-acetylaspartate reduction. NAC, but not taurine, prevented the HFD-induced reduction of lactate and phosphocreatine-to-creatine ratio. Analysis of the metabolite profile further revealed prominent NAC/taurine-induced increase of glutamate and GABA levels in the hippocampus. We conclude that NAC and taurine prevent of neuronal dysfunction, while only NAC prevents energy metabolism dysregulation in HFD-induced obesity in female mice.

## Introduction

Dementia represents a group of disorders characterized by cognitive decline that affects daily living activities, and it’s its prevalence is expected to nearly triple in the next 30 years, likely due to increased longevity and unhealthy lifestyles (GBD 2019 Dementia Forecasting Collaborators, 2022). Obesity has been proposed as a risk factor for developing dementia (Qizilbash *et al*., 2015; Albanese *et al*., 2017), since it is associated to hypertension, cardiovascular disease, metabolic syndrome and type 2 diabetes (T2D), which are modifiable risk factors for dementia (Livingston *et al*., 2020). Moreover, these vascular and metabolic factors might modulate genetic susceptibility for neurodegenerative disorders, namely Alzheimer’s disease (Guerreiro *et al.*, 2012).

Obesogenic diets rich in fat and sugar are commonly employed to experimentally induce metabolic syndrome that progresses into T2D. Rodents fed obesogenic diets display brain metabolism alterations (reviewed in García-Serrano & Duarte, 2020). By measuring metabolite profiles *in vivo* using magnetic resonance spectroscopy (MRS), we found that the brain of mice fed a high-fat diet (HFD) for 6 months shows increased concentrations of taurine, which was particularly prominent in the hippocampus (Lizarbe, Soares *et al*., 2019), and was also reported in the hippocampus of non-obese T2D Goto-Kakizaki rats (Duarte *et al*., 2019) and streptozotocin-induced type 1 diabetic rats (Duarte *et al*., 2009; Zhang *et al*., 2015). Recently, we measured metabolite alterations in the hippocampus and cortex over a 6-months period of high-fat and high-sucrose diet (HFHSD) feeding in mice (García-Serrano *et al*., 2022). Among the MRS-measured metabolites, concentrations of taurine increased in the hippocampus (but not in the cortex) at 8 weeks of HFHSD exposure, remained elevated for at least 4 months, and could be recovered to control levels by diet normalization. Interestingly, while the accumulation of taurine in the hippocampus of HFHSD-fed mice only appeared after glucose intolerance and insulin resistance were installed (García-Serrano *et al*., 2022), memory impairment has been reported within one week of HFD feeding (McLean *et al*., 2019; de Paula *et al*., 2021). Altogether, these findings suggest that brain taurine accumulation is unlikely to be related to the development of memory impairment in obesity and diabetes models, and might be a consequence of metabolic syndrome severity.

In contrast to these observations in rodent models of obesity and diabetes, mouse models for Alzheimer’s disease show reduced taurine concentration in the hippocampus and cortex, relative to controls (*e.g.* Aytan *et al*., 2016; Takado *et al*., 2018; Chiquita *et al*. 2019; Tondo *et al*., 2020). Interestingly, Takado and colleagues demonstrated an inverse relationship between absolute taurine concentration and tau protein accumulation in rTg4510 mice (Takado *et al*., 2018), and Aytan and colleagues showed an inverse correlation between the ratio of taurine-to-creatine and a brain GFAP levels in the 5xFAD model (Aytan *et al.*, 2016), suggesting that taurine depletion is associated with features of neurodegeneration and astrogliosis. Furthermore, taurine administration has been proposed to prevent neurodegeneration and improve memory performance (Louzada *et al*., 2004; Kim *et al.*, 2014).

Given this inverse relation between taurine levels and neurodegeneration, and the reported benefits of taurine administration, we propose that the accumulation of taurine in the hippocampus of obesity and diabetes models is a counteracting beneficial response to metabolic syndrome aggravation. Thus, the present study aimed at testing whether dietary taurine supplementation prevents HFD-induced memory impairment and metabolite profile alterations in the hippocampus. *N*-acetylcysteine (NAC) can act as a donor of cysteine for synthesis of taurine as well as glutathione, and NAC-treated new-born mice show a transient increase in cortical taurine levels (Duarte, Kulak *et al*., 2012). While, main biological effects of NAC are through increasing cysteine availability, it can also act as antioxidant, and interacts with dissulfide bonds and thus modulate thiolated proteins (Aldini *et al*., 2018). Accordingly, NAC is a well-tolerated drug that is suggested to exert beneficial effects on certain neurological and psychiatric disorders (Deepmala *et al*., 2015). Thus, we tested putative beneficial effects of stimulating of endogenous taurine and glutathione synthesis by NAC treatment, in parallel with taurine supplementation.

## Materials al Methods

### Animals

Animal experiments were performed according to EU Directive 2010/63/EU under approval of the Malmö/Lund Committee for Animal Experiment Ethics (permit number 5123/2021), and are reported following the ARRIVE guidelines (Animal Research: Reporting In Vivo Experiments, NC3Rs initiative, UK). Sample size was estimated from previous MRS experiments (García-Serrano *et al*., 2022). Female C57BL/6J mice (8-weeks old) were purchased from Taconic Biosciences (Köln, Germany). Mice were housed in groups of 5 on a 12h light-dark cycle with lights on at 07:00, room temperature of 21-23 °C and humidity at 55-60%. Mice were habituated to the facility for 2 weeks upon arrival. Mice were randomly assigned to 6 experimental groups (n=10/group): control diet (CD), a HFD, CD and HFD supplemented with 3%(w/v) taurine (>=98%, #W381306, Sigma-Aldrich, Germany) in drinking water (CD-T or HFD-T), and CD and HFD supplemented with 3%(w/v) *N*-acetylcysteine (99.9%, #A-2805, AG Scientific, San Diego, CA-USA) in the drinking water (CD-NAC or HFD-NAC). Dietary intervention and taurine/NAC supplementation started at 10 weeks of age and during 2 months (<u>figure 1A</u>). Food and water were provided *ad libitum*. Diets were acquired from Research Diets (New Brunswick, NJ-USA): a lard-based diet with 60% kcal of fat (D12492) and a control diet containing 10% kcal of fat (D12450J), with total energy of 5.21 and 3.82 kcal/g, respectively, and caloric intake was monitored as previously (García-Serrano *et al*, 2022).

**Figure 1.**
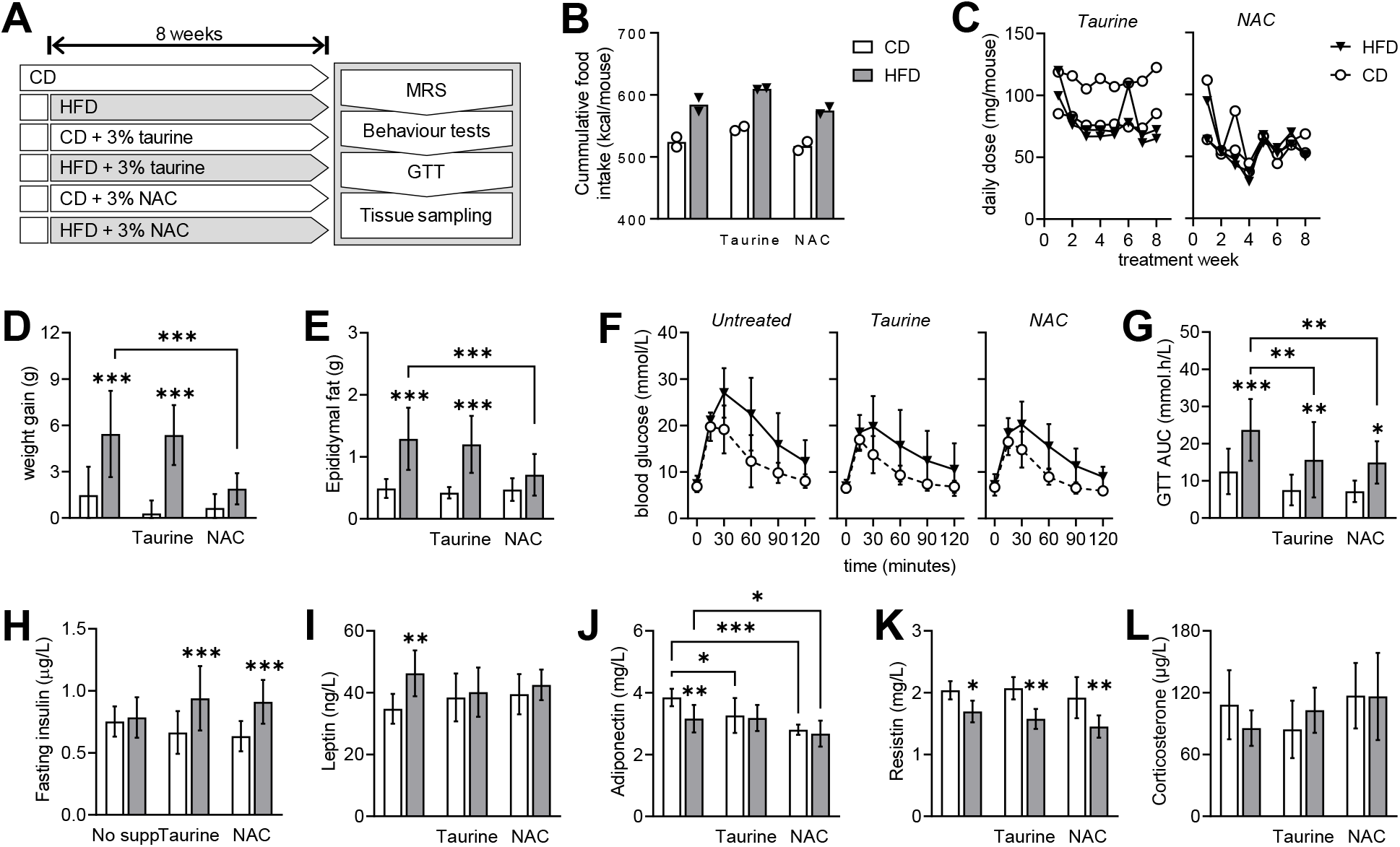
Study design and metabolic phenotype of female mice exposed to HFD and treated with either taurine or NAC. (A) After 8 weeks of treatment, all mice went through a behavioural testing, MRS and GTT before tissue collection. (B) Cages with HFD fed mice showed higher caloric intake than controls. (C) Estimated mean daily doses of taurine and NAC across the 8 treatment weeks was 85±20 and 58±15 mg/mouse. (D) Weight gain during treatment and (E) epididymal fat weight was larger in HFD-fed mice than controls, and prevented by NAC but not taurine treatment. (F) Glucose clearance in GTT was reduced by HFD feeding. (G) The area under the curve (AUC) of the GTT shows some effect of taurine and NAC on Glucose clearance. (H) Fasting insulin concentration in plasma was not affected by HFD, but was increased in HFD-fed mice treated with both taurine and NAC. Fed plasma concentrations of leptin (I), adiponectin (J), resistin (K) and corticosterone (L) were also impacted by either HFD or treatments. Data are shown as individual values of each cage in B-C, and as mean±SD of 10 mice in the remaining graphs. Open and filled bars/symbols represent CD- and HFD-fed mice, respectively. * P< 0.05, ** P< 0.01, *** P< 0.001 for comparison between HFD and CD within each treatment or as indicated, based on Fischer’s LSD post-hoc testing following a significant ANOVA effect of diet, supplementation or interaction.

### Magnetic Resonance Spectroscopy (MRS)

MRS was performed at 8 weeks under isoflurane anaesthesia as detailed in Garcia-Serrano *et al*. (2022). In brief, isoflurane (Vetflurane, Virbac, Carros, France) anaesthesia was delivered using 1:1 (v/v) O_2_:air as carrier gas at variable rate of 1-2% for maintaining stable respiration between 60 and 100 breaths per minute. Warm water circulation was used to keep body temperature at 36-37°C. MRS was acquired on a preclinical 9.4 T Bruker BioSpec AV III (Bruker, Ettlingen, Germany) equipped with a ^1^H quadrature transmit/receive cryoprobe, using STimulated Echo Acquisition Mode (STEAM) with repetition time of 4 s, echo time of 3 ms, mixing time of 20 ms, and spectral width of 4401.41 Hz. Water-supressed spectra were acquired in 20 blocks of 16 scans from a volume of interest (VOI) placed in the dorsal hippocampus (1.8 mm x 1.2 mm x 1.5 mm). An unsuppressed water spectrum from the same VOI was acquired in one block of 16 averages. After block alignment and summation in MATLAB (MathWorks, Natick, MA), metabolite concentrations were determined with LCModel v.6.3-1A (Stephen Provencher Inc., Oakville, Ontario-Canada; RRID: SCR_014455), including a macromolecule (Mac) spectrum in the database and using the unsuppressed water signal as internal reference (Duarte *et al*., 2014). The following metabolites were included in the LCModel analysis: alanine (Ala), ascorbate (Asc), aspartate (Asp), β-hydroxybutyrate, creatine (Cr), γ-aminobutyrate (GABA), glutamine (Gln), glutamate (Glu), glutathione (GSH), glycine (Gly), glycerophosphorylcholine (GPC), glucose (Glc), lactate (Lac), *myo*-inositol (Ins), *N*-acetylaspartate (NAA), phosphorylethanolamine (PE), phosphorylcholine (PCho), phosphocreatine (PCr), *scyllo*-inositol, and taurine (Tau). The Cramér-Rao lower bound (CRLB) was provided by LCModel as a measure of the reliability of the quantification for each metabolite, and metabolites with CRLB larger than 30% were disregarded as they did not fulfil reliability criteria. Namely, β-hydroxybutyrate, *N*-acetylaspartylglutamate, glucose and *scyllo*-inositol were not further analysed, and phosphorylcholine and glycerophosphorylcholine were analysed as total choline (PCho+GPC).

### Behaviour analyses

Experiments were performed from 9:00 to 18:00, with light adjusted to an illuminance of 15 lx in each apparatus. After acclimatizing to the testing room for at least 1 hour, open field test and object recognition tasks for testing novel object recognition (NOR) or novel location recognition (NLR) were conducted as detailed in Garcia-Serrano *et al*. (2022). Exploration was observed in a cubic arena with a side length of 50 cm. Mice were first habituated to the empty arena for 8 minutes. Arena exploration was analysed with Any-maze behavioural tracking software (version 6.0., Dublin, Ireland). Total walk distance, number of crossings between arena quadrants and immobility time, as well as exploration of the arena center at 6 cm from the walls were tracked. Thereafter, NLR was assessed by placing the mice in the arena with two identical objects, and allowed to explore them for 5 minutes (familiarization phase). Mice were then removed from the arena for 1 hour (retention phase), and reintroduced for 5 minutes but with one object relocated to a different quadrant in the arena (recognition phase). For NOR, two new identical objects were used in the familiarization phase, and one of them was replaced by a novel object during the recognition phase. Time exploring each object was measured.

### Glucose tolerance test (GTT)

Food was removed for 6 hours starting at 08:00. After, a blood sample was collected from the vena saphena to determine plasma insulin levels by ELISA (#10-1247-10, Mercodia, Uppsala, Sweden; RRID: AB_2783837). Blood glucose was measured from tail tip blood with the Breeze glucometer (Bayer, Zürich, Switzerland). Then mice were given 2 g/kg glucose i.p. from a 30%(w/v) solution in saline, followed by determination of glucose levels at 15, 30, 60, 90, and 120 minutes.

### Tissue collection

At the end of the study, mice were anesthetized with isoflurane, and blood was drawn from the portal vein into heparinized tubes kept on ice. Mice were decapitated after cardiac perfusion with 20 mL of cold phosphate-buffered saline (PBS; in mmol/L: 137 NaCl, 2.7 KCl, 1.5 KH_2_PO_4_, 8.1 Na_2_HPO_4_, pH 7.4). The hippocampi were dissected and quickly frozen in N2 (l), and then stored at - 80 °C. Collected blood samples were centrifuged at 21,000×g and 4 °C for 15 minutes, and plasma was frozen at −80 °C.

### Plasma hormones

ELISA kits from Abcam (Cambridge, UK) were used to determine plasma leptin (#ab100718, RRID: AB_2889903), adiponectin (#ab108785, AB_2891131), resistin (#ab205574, AB_2891132) and corticosterone (#ab108821, AB_2889904) in samples collected upon mouse sacrifice.

### Plasma taurine determination

Plasma metabolites were extracted by mixing 300 μL of methanol with 70 μL of plasma, followed by sonication on ice for 30 minutes, and centrifugation at 13,000×g and 4 °C for 30 minutes. Supernatants were dried using a Savant SpeedVac Concentrator, and then ressuspended in 100 mM sodium phosphate buffer pH 7.4 prepared in ^2^H_2_O (>99.9%, Sigma-Aldrich), containing 0.01% NaN_3_. Sodium fumarate (5 μmol/L) was included as internal standard for quantification by ^1^H nuclear magnetic resonance (NMR) spectroscopy (Duarte *et al*., 2007). Spectra were acquired at 25 °C on an 11.7 T Varian Inova spectrometer equipped with a 5 mm broadband probe (Agilent technologies, Santa Clara, CA, USA), and using a Carr-Purcell-Meiboom-Gill sequence with spectral width of 8,012.8 Hz, acquisition time of 2.045 s, and relaxation delay of 10 s. All spectra were collected with 520 acquisitions. Taurine concentration was determined by measuring peak areas using NUTS (Acorn NMR, Fremont, USA).

### Real-time polymerase chain reaction (RT-PCR)

RNA was isolated from the hippocampus with Trizol (#15596026, Invitrogen), and then 1 μg of total RNA was reverse transcribed with random hexamer primers using the qScript cDNA SuperMix (#95048, Quantabio, England), according to the manufacturers’ instructions. The resulting cDNA was used for quantitative RT-PCR as detailed previously (Garcia-Serrano *et al*., 2022) using PerfeCTa SYBRgreen SuperMix (#95054, Quantabio, England) and the primers listed in <u>table 1</u>. Gene expression was determined using the comparative cycle threshold (CT) method (ΔΔCT) with normalization to the average expression of L14 and GAPDH.

**Table 1.**
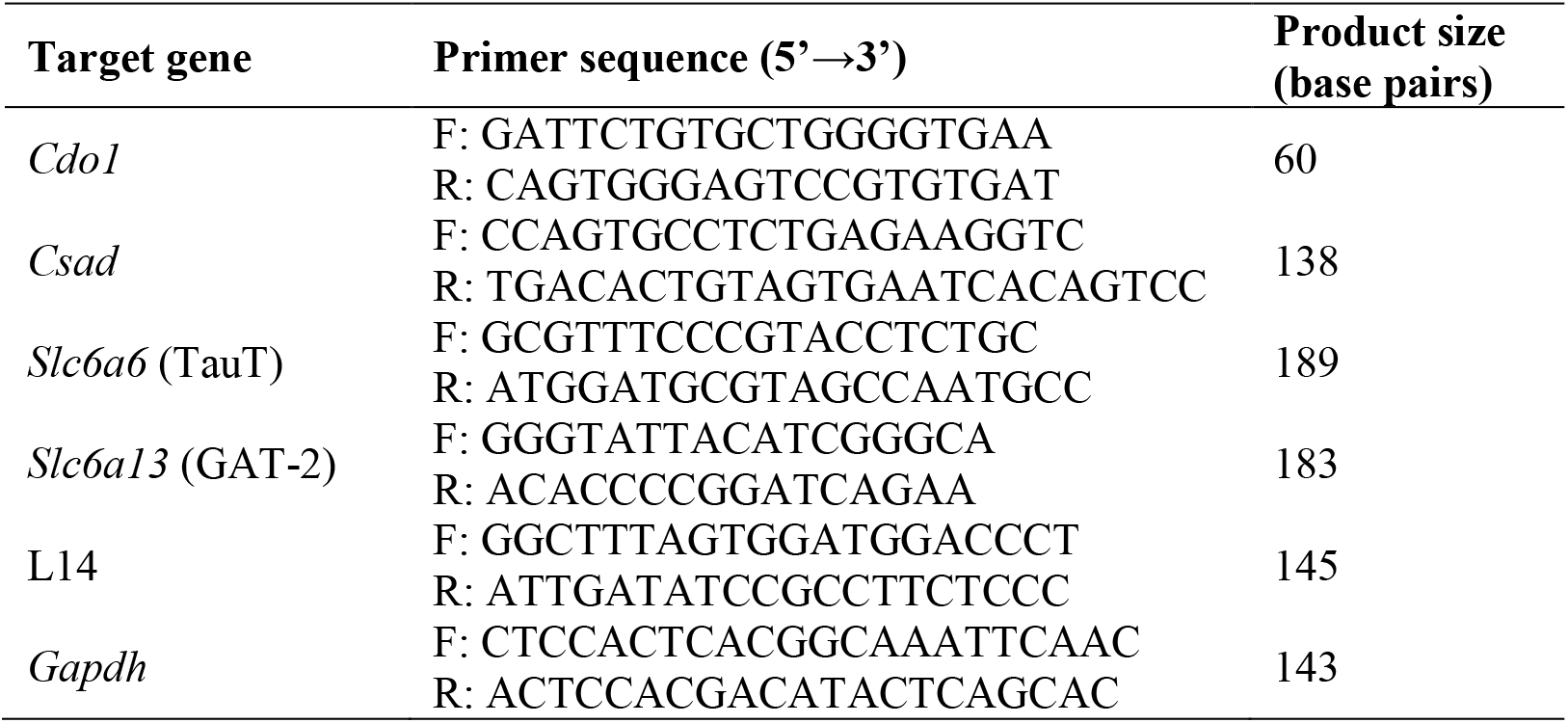
Sequence of primers used for RT-PCR and respective product size.

### Immunoblotting

Western blotting of hippocampal protein extracts was carried out as detailed previously (Lizarbe *et al*., 2019) with antibodies against the taurine transporter TauT (SLC6a6, #PA5-37460, Invitrogen, Life Technologies Europe, Stockholm, Sweden; RRID: AB_2554074), GABA transporter GAT-2 (SLC6a13, #PA5-113493, Invitrogen; RRID: AB_2868226) and GAPDH (#ab9485, Abcam; RRID: AB_307275).

### Statistical analysis

All statistical analyses were carried out in Prism 9.3.0 (GraphPad, San Diego, CA-US; RRID: SCR_002798). After confirming that data showed no serious deviation from normality, data were analysed by 2-way ANOVA with diet and treatment as factors. Upon significant effects of diet, supplementation or their interaction, the Fisher’s least significant difference (LSD) test was used for independent comparisons between HFD and control groups within each treatment, or between treated and untreated groups within each diet.

## Results

### Food intake and metabolic phenotype

Total caloric intake over 2 months was higher for cages with HFD-fed mice than controls, with minimal effects across treatments (<u>figure 1B</u>). Estimated daily doses of taurine and NAC were on average 78 (range 38-122) and 67 (range 30-120) mg/mouse (<u>figure 1C</u>). After 2 months, HFD-fed mice showed higher weight gain than controls, which was prevented by NAC but not taurine treatment (<u>figure 1D</u>; diet F(1,54)=60.42, P<0.0001; treatment F(2,54)=8.720, P=0.0005, interaction F(2,54)=6.688, P=0.0025). Similarly, when compared to the respective controls, epididymal fat was increased in HFD and taurine-treated HFD but not in NAC-treated HFD (<u>figure 1E</u>; diet F(1,54)=51.72, P<0.0001; treatment F(2,54)=4.522, P=0.0153, interaction F(2,54)=4.728, P=0.0128).

Glycaemia after a 6-hour fasting was similar across all groups (<u>figure 1F</u>; diet F(1,54)=0.7442, P=0.3922; treatment F(2,54)=0.8636, P=0.4275, interaction F(2,54)=0.5180, P=0.5987). Upon glucose administration, HFD-fed mice in any group showed slower glucose clearance than the respective controls (<u>figure 1F</u>). However, the area under the curve of the glucose tolerance test indicates that both taurine and NAC improved glucose clearance in HFD-fed mice (<u>figure 1G</u>; diet F(1,54)=27.45, P<0.0001; treatment F(2,54)=6.920, P=0.0021, interaction F(2,54)=0.3920, P=0.6776).

In contrast to males, female mice are resistant to develop insulin resistance upon exposure to obesogenic diets (Garcia-Serrano *et al*., 2022). Accordingly, HFD did not elicit fasting hyperinsulinaemia, but treatments with taurine and NAC caused a small increase in circulating insulin in HFD-fed but not CD-fed mice (<u>figure 1H</u>; diet F(1,54)=18.55, P<0.0001; treatment F(2,54)=0.2112, P=0.8103, interaction F(2,54)=3.225, P=0.0475). We further analysed levels of circulating hormones that act on the brain, namely adipokines and corticosterone. Leptin was increased in HFD-fed mice *versus* controls but not upon Taurine or NAC treatment (<u>figure 1I</u>; diet F(1,54)=5.860, P=0.0217; treatment F(2,54)=0.2043, P=0.8163, interaction F(2,54)=1.878, P=0.1705). Adiponectin was reduced by HFD-feeding and further reduced by Taurine and NAC treatments (<u>figure 1J</u>; diet F(1,54)=4.876, P=0.0350; treatment F(2,54)=11.01, P=0.0003, interaction F(2,54)=2.098, P=0.1404). Resistin was lowered by HFD-feeding with no effect treatments (figure 1K; diet F(1,54)=38.56, P<0.0001; treatment F(2,54)=2.462, P=0.1029, interaction F(2,54)=0.4405, P=0.6479). Corticosterone levels showed no effects of diet or treatment (figure 1L; diet F(1,54)=0.02658, P=0.8716; treatment F(2,54)=2.038, P=0.1479, interaction F(2,54)=1.401, P=0.2620).

### Taurine homeostasis

Mice were treated with either taurine or NAC that acts as a cysteine donor for taurine synthesis through oxygenation and decarboxylation reactions catalysed by cysteine dioxygenase 1 (CDO1) and cysteine sulfinic acid decarboxylase (CSAD) (<u>figure 2A</u>). Brain taurine is not only synthesized locally, it is also transported from the periphery via the taurine transporter TauT (*Slc6a6*) and the GABA transporter GAT-2 (*Slc6a13*) (Zhou *et al*., 2012). HFD had no significant impact on the relative hippocampal expression of these four players in taurine homeostasis (<u>figure 2B</u>). NAC treatment decreased CSAD expression in CD-fed mice (P=0.0421 *vs*. untreated CD), and no effects of NAC were observed (<u>figure 2B</u>; diet F(1,38)=4.450, P=0.0415; treatment F(2,38)=1.742, P=0.1889, interaction F(2,38)=0.5822, P=0.5636). Western blot analysis of TauT and GAT-2 did not reveal significantly altered density of either taurine carrier in the hippocampus (figure 2C). NMR spectroscopy analysis of plasma extracts showed that treatment with taurine increases its concentration in circulation (figure 2D; diet F(1,40)=1.088, P=0.3031; treatment F(2,40)=7.366, P=0.0019, interaction F(2,40)=1.272, P=0.2914). Surprisingly, NAC had no effect on plasma taurine concentration. Similar alterations were observed for taurine concentration in the hippocampus (see below).

**Figure 2.**
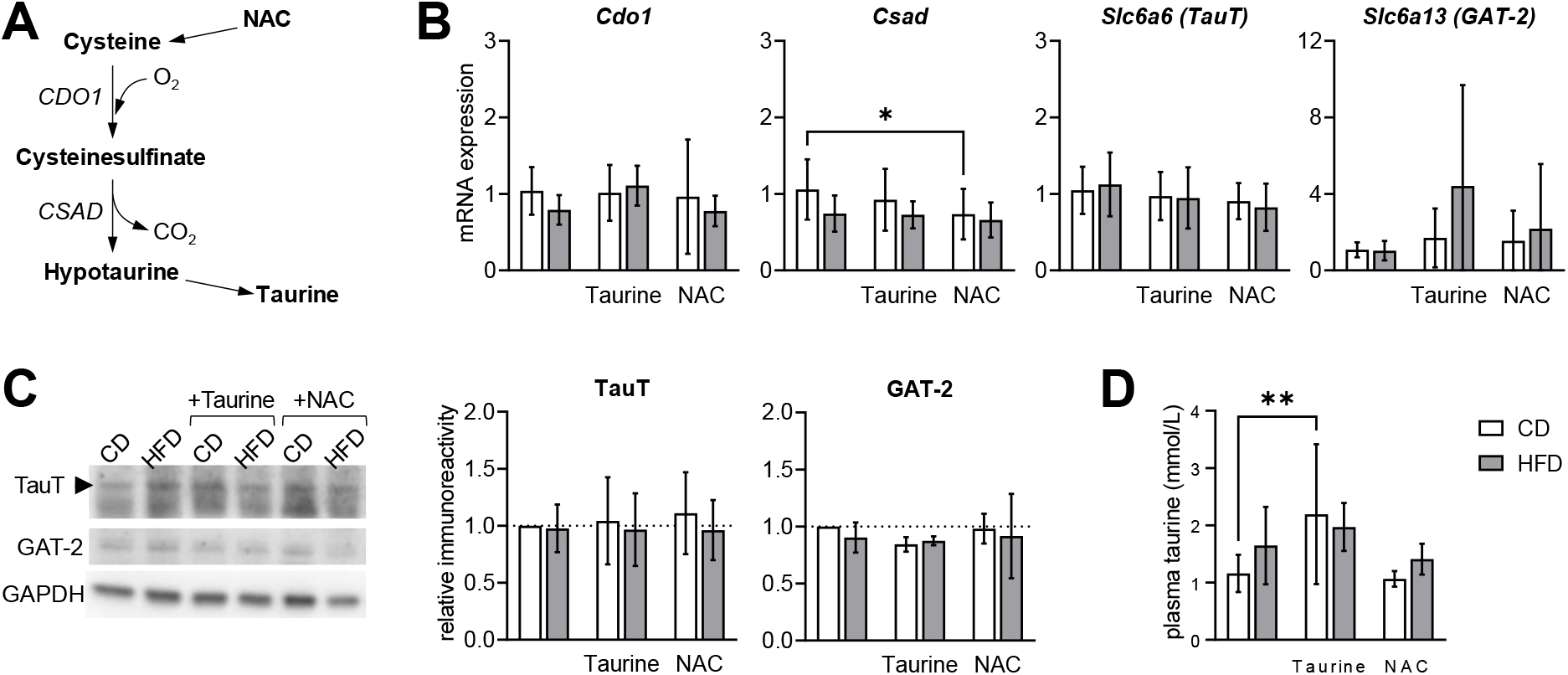
Taurine homeostasis in mice treated with taurine and NAC. (A) Schematic representation of endogenous synthesis of taurine from cysteine of NAC origin through oxygenation and decarboxylation reactions catalysed by cysteine dioxygenase 1 (CDO1) and cysteine sulfinic acid decarboxylase (CSAD). (B) Relative mRNA expression of *Cdo1*, *Csad* and taurine carriers TauT (*Slc6a6*) and GAT-2 (*Slc6a13*) in the hippocampus. (C) Western blot analysis of hippocampal TauT and GAT-2 density. (D) Concentration of taurine in plasma. Data are shown as mean±SD of n=6-8 in B, n=4 in C, n=6-10 in D. Open and filled bars/symbols represent CD- and HFD-fed mice, respectively. * P< 0.05, ** P< 0.01, *** P< 0.001 based on Fischer’s LSD post-hoc comparison after significant effect of diet, supplementation or interaction in ANOVA.

### Taurine and NAC treatments prevent HFD-induced memory impairment

Behaviour was assessed in object exploration tasks. Mice were firstly allowed to freely explore the empty area. Such open-field exploration was not impacted by HFD exposure or treatments with taurine or NAC, as revealed by similar total walking distance, number of crossings between quadrants, total immobility time, fraction of time spent in arena center (<u>figure 3A</u>). These results suggest the absence of stress or anxiety-like behaviour induced by any treatment. After, memory performance was evaluated with the novel object recognition (NOR) and novel location recognition (NLR) tests. Relative to controls, untreated HFD-fed mice showed impaired memory as depicted by reduced preference for novelty in NOR and NRL (respectively, P=0.0250 and P=0.0414 *vs*. CD). On the other hand, memory performance was preserved in HFD-fed mice treated with either taurine or NAC in the NOR task (<u>figure 3B</u>; diet F(1,53)=4.947, P=0.0304; treatment F(2,53)=0.5773, P=0.5649, interaction F(2,53)=0.8513, P=0.4326), as well as in the NLR task (<u>figure 3C</u>; diet F(1,53)=4.780, P=0.0332; treatment F(2,53)=2.716, P=0.0754, interaction F(2,53)=0.5347, P=0.5890).

**Figure 3.**
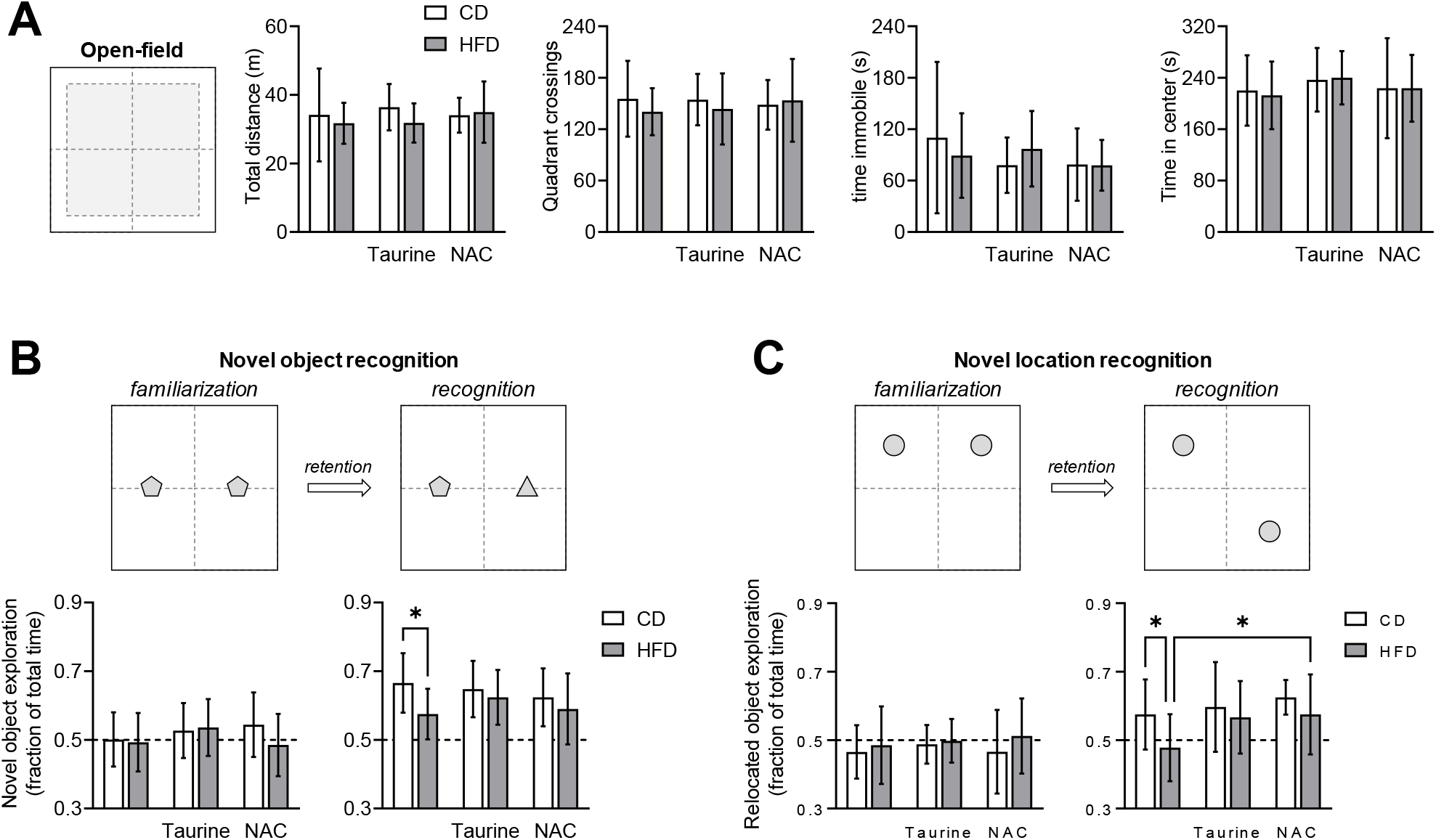
HFD-induced memory impairment was prevented by taurine and NAC treatments. (A) Open-field exploration was not impacted by HFD exposure or treatments with taurine or NAC, as revealed by similar total walking distance, number of crossings between quadrants, total immobility time, fraction of time spent in arena center. (B-C) Novel object recognition (NOR) and novel location recognition (NLR) tests showed identical object exploration in the familiarization phase for all groups (mean at ~50% for each object; graphs on the left)), while impaired novelty recognition was identified by both NOR (B) and NLR (C) in untreated HFD-fed mice. Open and filled bars represent mice fed CD and HFD, respectively. Data are expressed as mean±SD of generally n=10. Due to object exploration below 10 seconds, one non-supplemented CD-fed mouse was excluded from NOR in recognition phase (n=9), and one taurine-supplemented HFD-fed mouse was excluded from NLR in recognition phase (n=9). * P< 0.05 based on Fischer’s LSD post-hoc comparison after significant effect of diet, supplementation or interaction in ANOVA.

### Metabolite profile alterations in the hippocampus

Metabolic alterations induced by long-term diet-induced obesity occur throughout the whole brain, but are particularly prominent in the hippocampus (Lizarbe *et al*., 2019; Garcia-Serrano *et al*., 2022). Therefore, we set to measure the metabolite profile in this brain area (<u>figure 4; table 2</u>). As expected, supplementation with taurine increased its concentration in the hippocampus of mice fed either CD of HFD (P<0.001 for both relative to the respective untreated group; <u>figure 4A</u>). NAC treatment had a smaller effect on hippocampal taurine, which was only significantly increased in HFD fed mice (P=0.0268 *vs*. untreated HFD). *N-*acetylaspartate concentration, which is considered a marker or neuronal health (Duarte, Lei *et al*., 2012), was decreased by HFD consumption (P=0.0071 *vs.* CD), but not upon supplementation with either taurine or NAC (<u>Figure 4A</u>). Lactate concentration was also decreased by HFD feeding (P=0.0045 *vs.* CD), but not upon NAC treatment (<u>Figure 4A</u>). A similar trend was observed for hippocampal alanine levels. Small variations in phosphocreatine and creatine were observed in the hippocampus of HFD-fed mice (<u>figure 4B</u>). Most importantly, their ratio was significantly reduced by HFD (P=0.0139 *vs*. CD), a modification that was prevented by treatment with NAC but not taurine (<u>figure 4B</u>; diet F(1,54)=9.753, P=0.0029; treatment F(2,54)=2.040, P=0.1400, interaction F(2,54)=0.8315, P=0.4409).

**Table 2.**
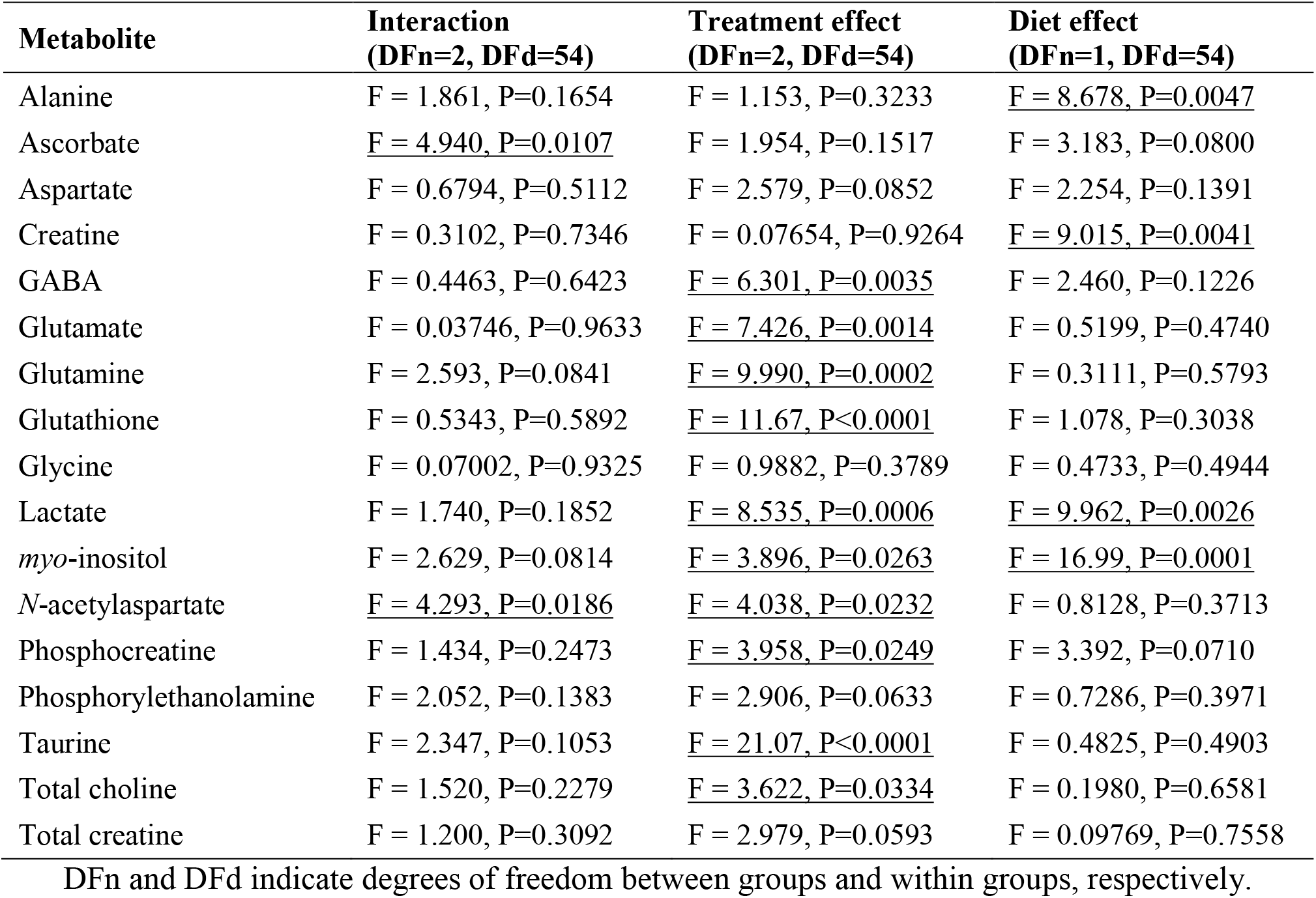
ANOVA results F and P values for the analyses of metabolite concentrations in the hippocampus (n=10/group).

| **Metabolite** | **Interaction**<br>**(DFn=2, DFd=54)** | **Treatment effect**<br>**(DFn=2, DFd=54)** | **Diet effect**<br>**(DFn=1, DFd=54)** |
| --- | --- | --- | --- |
| Alanine | F = 1.861, P=0.1654 | F = 1.153, P=0.3233 | F = 8.678, P=0.0047 |
| Ascorbate | <u>F = 4.940, P=0.0107</u> | F = 1.954, P=0.1517 | F = 3.183, P=0.0800 |
| Aspartate | F = 0.6794, P=0.5112 | F = 2.579, P=0.0852 | F = 2.254, P=0.1391 |
| Creatine | F = 0.3102, P=0.7346 | F = 0.07654, P=0.9264 | <u>F = 9.015, P=0.0041</u> |
| GABA | F = 0.4463, P=0.6423 | <u>F = 6.301, P=0.0035</u> | F = 2.460, P=0.1226 |
| Glutamate | F = 0.03746, P=0.9633 | <u>F = 7.426, P=0.0014</u> | F = 0.5199, P=0.4740 |
| Glutamine | F = 2.593, P=0.0841 | <u>F = 9.990, P=0.0002</u> | F = 0.3111, P=0.5793 |
| Glutathione | F = 0.5343, P=0.5892 | <u>F = 11.67, P<0.0001</u> | F = 1.078, P=0.3038 |
| Glycine | F = 0.07002, P=0.9325 | F = 0.9882, P=0.3789 | F = 0.4733, P=0.4944 |
| Lactate | F = 1.740, P=0.1852 | <u>F = 8.535, P=0.0006</u> | <u>F = 9.962, P=0.0026</u> |
| *myo*-inositol | F = 2.629, P=0.0814 | <u>F = 3.896, P=0.0263</u> | <u>F = 16.99, P=0.0001</u> |
| *N*-acetylaspartate | <u>F = 4.293, P=0.0186</u> | <u>F = 4.038, P=0.0232</u> | F = 0.8128, P=0.3713 |
| Phosphocreatine | F = 1.434, P=0.2473 | <u>F = 3.958, P=0.0249</u> | F = 3.392, P=0.0710 |
| Phosphorylethanolamine | F = 2.052, P=0.1383 | F = 2.906, P=0.0633 | F = 0.7286, P=0.3971 |
| Taurine | F = 2.347, P=0.1053 | <u>F = 21.07, P<0.0001</u> | F = 0.4825, P=0.4903 |
| Total choline | F = 1.520, P=0.2279 | <u>F = 3.622, P=0.0334</u> | F = 0.1980, P=0.6581 |
| Total creatine | F = 1.200, P=0.3092 | F = 2.979, P=0.0593 | F = 0.09769, P=0.7558 |

DFn and DFd indicate degrees of freedom between groups and within groups, respectively.

**Figure 4.**
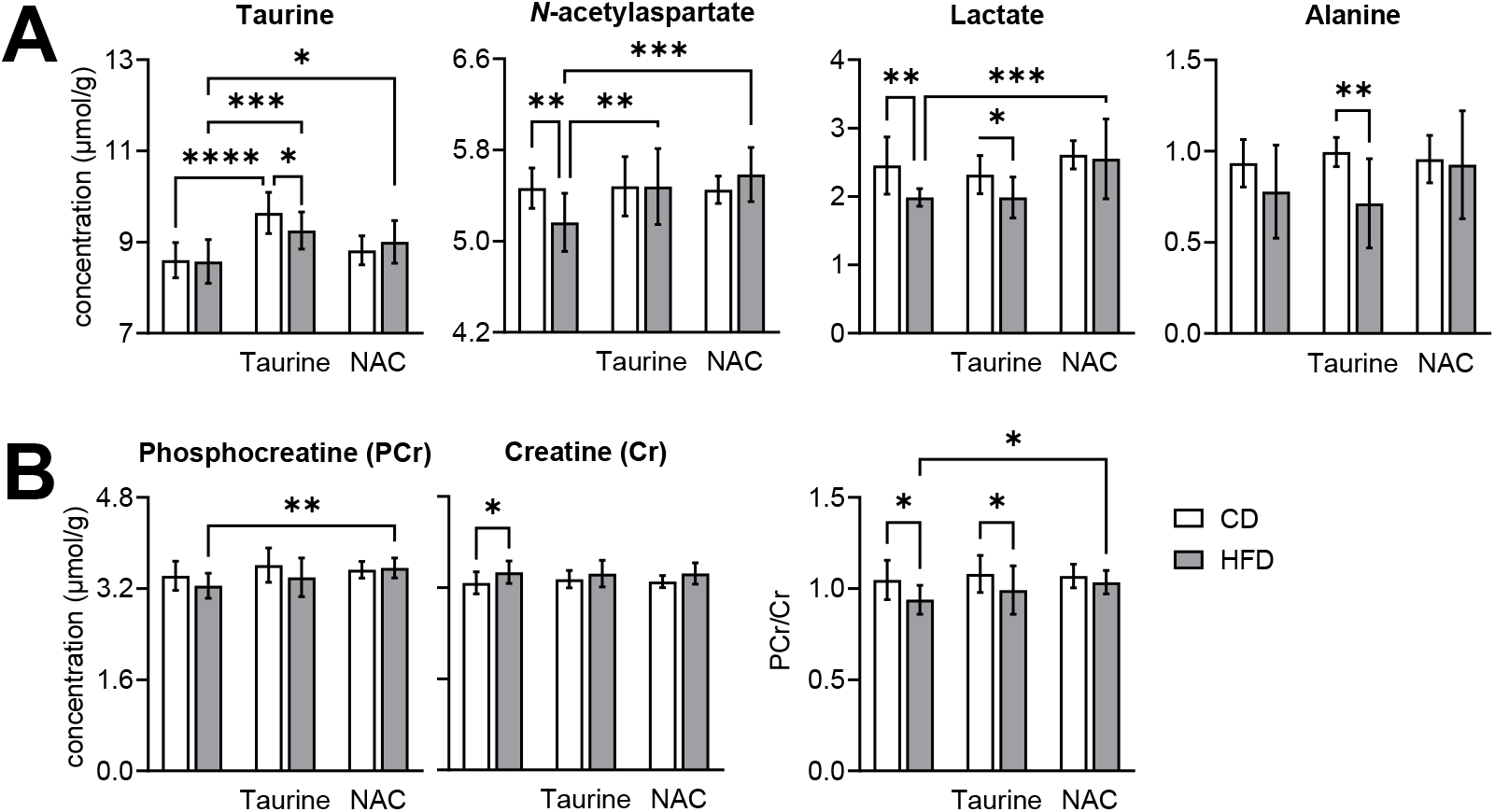
HFD-induced alterations of metabolite concentrations in the hippocampus. (A) Treatment with taurine and/or NAC increased hippocampal taurine concentration, and prevented the HFD-induced reduction in *N*-acetylaspartate and/or lactate levels. (B) NAC treatment prevented the HFD-induced reduction of phosphocreatine-to-creatine ratio. Open and filled bars represent mice fed CD and HFD, respectively. Data are expressed as mean±SD of n=10. * P< 0.05, ** P< 0.01, *** P< 0.001, **** P< 0.0001 based on Fischer’s LSD post-hoc comparison after significant effect of diet, supplementation or interaction in ANOVA.

A number of metabolites in the neurochemical profile were not modified by HFD exposure *per se*, but their concentrations in the hippocampus were increased by taurine and/or NAC treatment (<u>figure 5, table 2</u>). Both taurine and NAC increased the content of glutamate in both CD-fed and HFD-fed mice (all P<0.05 relative to the respective untreated group), While taurine had no impact on glutamine, NAC increased glutamine concentration in the hippocampus of HFD-fed mice (P<0.0001 *vs*. untreated HFD; P=0.0342 *vs*. NAC-treated CD). Similarly, NAC but not taurine treatment increased total choline concentration in HFD-fed mice (P=0.0061 *vs*. untreated HFD). Furthermore, in the hippocampus of both CD and HFD-fed mice, NAC increased the concentration of GABA (for both, P<0.05 compared to untreated groups) and glutathione (for both, P<0.01 compared to untreated groups). On the other hand, taurine treatment increased ascorbate concentration in CD-fed mice (P=0.0101 compared to untreated group) but not HFD-fed mice. Finally, while *myo*-inositol was not affected by treatments in CD-fed mice, *myo*-inositol concentration increased in the hippocampus of HFD-fed mice upon treatments with taurine (P=0.0291 *vs.* untreated HFD) and NAC (P=0.0019 *vs.* untreated HFD; P<0.0001 *vs*. NAC-treated CD).

**Figure 5.**
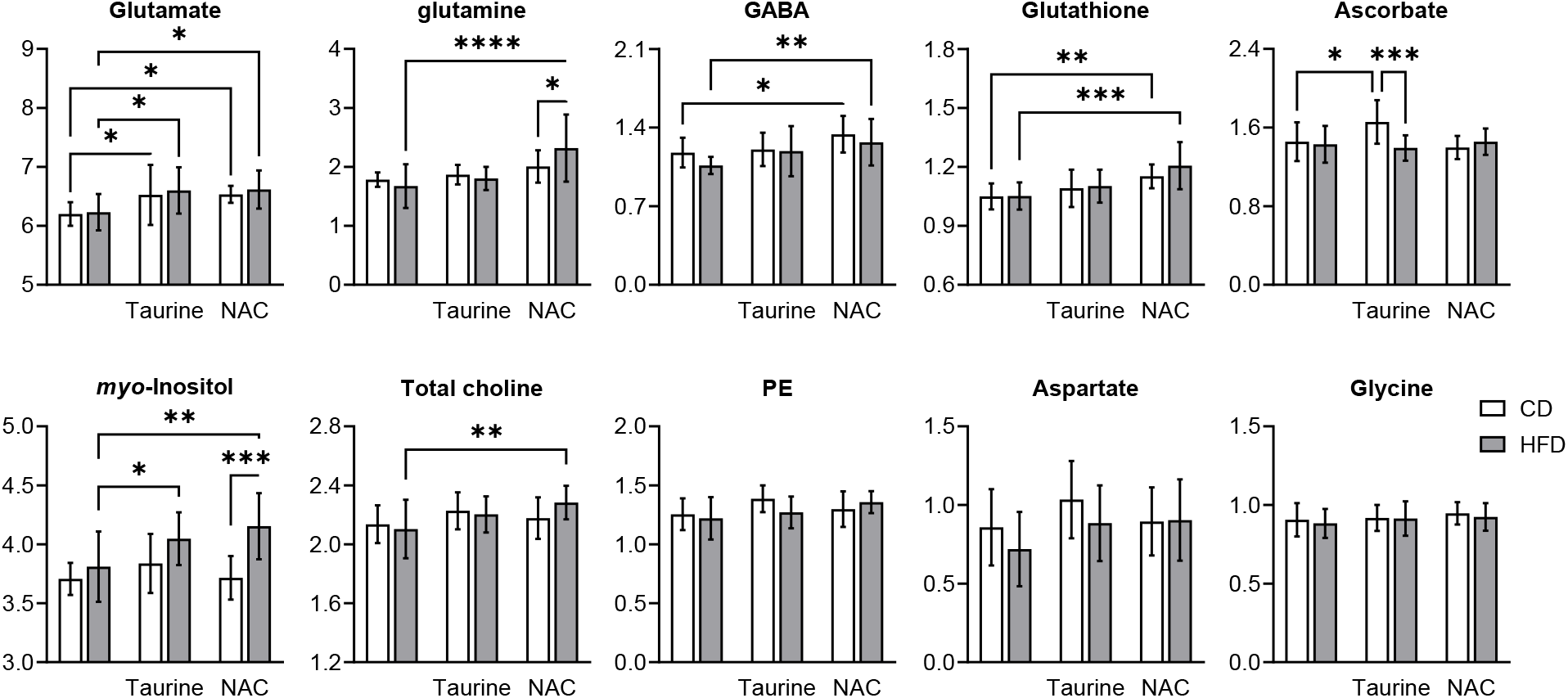
Taurine and NAC treatments impacted metabolite concentrations in the hippocampus. Open and filled bars represent mice fed CD and HFD, respectively. Data are expressed as mean±SD of n=10. * P< 0.05, ** P< 0.01, *** P< 0.001, **** P< 0.0001 based on Fischer’s LSD post-hoc comparison after significant effect of diet, supplementation or interaction in ANOVA.

## Discussion

This study demonstrates that taurine and NAC treatments have beneficial effects on HFD-induced alterations of hippocampal metabolism, and prevent the development of memory impairment in female mice fed a HFD for 2 months. Accordingly, taurine has also been proposed to improve memory in animal models such as the APP/PS1 mouse model of Alzheimer’s disease (Kim *et al*., 2014), aged mice (El Idrissi, 2008), or rats exposed to noise stress (Haider *et al*, 2020); and NAC was suggested to prevent memory impairment in an Alzheimer’s disease model of intra-hippocampal administration of amyloid-β (Shahidi *et al*., 2017) or amyloid-β oligomers (More *et al*., 2018).

HFD exposure for 2 months caused a marked reduction of *N-*acetylaspartate concentration, which is synthesized from mitochondrial acetyl-CoA and aspartate in neurons (Baslow, 2003), as thus it is generally taken as a biomarker of neuronal health (Duarte, Lei *et al*., 2012). Moreover, HFD triggered a reduction of the energy indicator phosphocreatine-to-creatine ratio, and the concentration of the glycolytic end-product lactate. Taken together, these three metabolic alterations suggest an overall depression of energy metabolism in the hippocampus, including deactivation of glycolysis and reduced mitochondrial activity, which may have a detrimental impact on neuronal function (Belenguer *et al*., 2019). Notably, NAC supplementation was able to prevent these alterations. On the other hand, taurine was only able to prevent the HFD-induced reduction of *N-*acetylaspartate levels.

We had hypothesized that NAC stimulates endogenous taurine since both neurons and astrocytes can produce taurine from cysteine (Brand *et al*., 1998; Vitvitsky *et al*., 2011). However, it is interesting to note that cultured astrocytes use exogenously cysteine to produce hypotaurine, taurine and glutathione at similar rates, neurons in culture produce glutathione at a much faster rate than any other metabolite (Vitvitsky *et al*., 2011). In our study, concentrations of taurine in both plasma (<u>figure 2D</u>) and hippocampus (<u>figure 4A</u>) increased with taurine supplementation but were unaltered by NAC treatment, suggesting that there is limited taurine synthesis from NAC-derived cysteine. Accordingly, CSAD expression was slightly reduced by NAC treatment. On the other hand, increased glutathione levels were observed in the hippocampus of NAC-treated mice, particularly prominent in HFD-fed mice (<u>figure 5</u>), which is in line with stimulation of glutathione synthesis. Thus, modes of action of the treatments are likely independent, with NAC effects being mediated via glutathione and not taurine metabolism. Through stimulation of the synthesis of the major antioxidant glutathione (via γ-glutamylcysteine ligase), NAC can prevent brain oxidative stress and neuroinflammation (*e.g*. Dwir *et al*., 2021). Accordingly, Choy *et al*. (2010) reduced glutathione brain levels in rats by treatment with 2-cyclohexene-1-one, and demonstrated that memory impairment is recoverable by NAC administration.

Further analysis of the metabolite profile of the hippocampus provides insight into putative mechanisms of action of taurine and NAC. Increased hippocampal taurine carries benefits *per se*, including promotion of neurogenesis and synaptogenesis (Shivaraj *et al*. 2012), which is in line with increased concentration of the neuronal marker *N*-acetylaspartate, as well as glutamate. Indeed, both taurine and NAC caused a robust accumulation of glutamate in the hippocampus. Glutamate is the main excitatory neurotransmitter, mainly located in neurons, and interfaces energy metabolism and neuronal activity (reviewed in Sonnay *et al*., 2017). Glutamate is key for memory performance, but excitatory glutamatergic neurons require adequate inhibitory control (Duarte & Xin, 2019). NAC but not taurine induced an accumulation of the inhibitory neurotransmitter GABA in the hippocampus. In this way, NAC can increase the inhibitory tonus. On the other hand, taurine itself is known to act as neurotransmitter via GABAA, GABAB and glycine receptors (Lynch *et al*., 2004; Albrecht & Schousboe, 2005). In line with a role of NAC treatment on neurotransmitter metabolism, glutamine concentration was also increased by NAC supplementation in the hippocampus of HFD mice. Given that glutamine is exclusively synthesized in glia (Sonnay *et al*., 2017), NAC treatment appears to be a modulator of the glutamate-glutamine cycle between neurons and astrocytes.

Dietary supplementation with either taurine or NAC increased the concentration of *myo*-inositol in the hippocampus of HFD-fed mice but not controls. In this study, HFD feeding alone did not impact hippocampal levels of *myo*-inositol. Increases in the content of *myo*-inositol are believed to represent astrogliosis (Duarte, Lei *et* al., 2012; Harris *et al*., 2015), but such relation has not been replicated in the hippocampus of diabetes models (Duarte, Carvalho *et al*., 2009; Duarte, Skoug *et al*., 2019). While the interpretation of *myo*-inositol changes remains unclear, one can speculate that taurine and NAC treatments are unlikely induce gliosis in the hippocampus. Similarly, the NAC-induced increase in total choline in HFD-fed mice relative to untreated HFD-fed mice is unlikely linked to neuroinflammation as proposed in other studies (see Yasmin *et al*., 2019), and simply represents the abundance of mobile metabolites resulting from enzymatic modification of phosphatidylcholine in cell membranes unrelated to gliosis (Iorio *et al*., 2021).

Taurine and NAC supplementation not only impacted the brain, they also ameliorated metabolic syndrome upon HFD feeding. Female mice under HFD developed obesity and glucose intolerance but not substantial hyperglycemia or hyperinsulinemia, as observed upon HFHSD feeding (Garcia-Serrano *et al*., 2022). NAC but not taurine prevented obesity development. As observed by others, HFD-induced glucose intolerance was prevented by treatment with taurine (Ribeiro *et al*., 2012; Camargo *et al*. 2013; Figueroa *et al*., 2016) and NAC (Falach-Malik *et al*. 2016). Both treatments modulated circulating levels of adipokines, in line with possible effects of NAC (*e.g.* Shen *et al*., 2018) and taurine (*e.g.* Kim *et al*., 2019) on adipose tissue.

The present study was limited to female mice. While most studies have been conducted on male rodents, our recent work suggests that the development of memory impairment promoted by HFHSD is similar in mice of either sex despite higher metabolic syndrome severity in males (García-Serrano *et al*, 2022). In particular, in contrast to males, female mice under HFHSD did not develop hyperinsulinemia. By focusing on female mice in the present work, we were able to study obesity-associated effects in the absence of hyperinsulinemia, and thus increased insulin concentration in the brain.

In conclusion, memory impairment and metabolic alterations induced by HFD on the hippocampus of female mice are partially prevented by taurine supplementation, and fully recovered by treatment with NAC. NAC acts as a cysteine donor that could stimulate taurine synthesis, but it mainly resulted in accumulation of glutathione in the hippocampus, likely providing an enhanced antioxidant defence. Mechanisms by which these treatments exert neuroprotection and prevent memory impairments involve energy metabolism, and the synthesis of glutamate and/or GABA.

## Abbreviations

AD: Alzheimer’s disease
Aβ: amyloid β
CD: control diet
CDO1: cysteine dioxygenase 1
CSAD: cysteine sulfinic acid decarboxylase
GAT-2: GABA transporter 2
HFD: high-fat diet
HFHSD: high-fat and high-sucrose diet
MRS: magnetic resonance spectroscopy
NAC: *N*-acetylcysteine
NLR: novel location recognition
NMR: Nuclear magnetic resonance
NOR: novel object recognition
PBS: phosphate-buffered saline
T2D: type 2 diabetes
TauT: taurine transporter
VOI: volume of interest

## Data availability statement

All data are contained within the manuscript, and can be shared upon request to the corresponding author.

## Acknowledgements

The authors thank Dr. Vladimir Denisov for access to NMR spectrometer. The Lund University Bioimaging Centre is acknowledged for providing access to 9.4T MRI scanner.

## Authors’ contributions

JMND designed the study and analysed data. AMGS, JPPV and VF performed experiments and analysed data. JMND and AMGS wrote the manuscript. All authors revised the manuscript.

## References

Albanese E, Launer LJ, Egger M, Prince MJ, Giannakopoulos P, Wolters FJ, Egan K. Body mass index in midlife and dementia: Systematic review and meta-regression analysis of 589,649 men and women followed in longitudinal studies. Alzheimers Dement (Amst). 2017 Jun 20;8:165–178. doi: 10.1016/j.dadm.2017.05.007.

Albrecht J, Schousboe A. Taurine interaction with neurotransmitter receptors in the CNS: an update. Neurochem Res. 2005 Dec;30(12):1615–21. doi: 10.1007/s11064-005-8986-6.

Aldini G, Altomare A, Baron G, Vistoli G, Carini M, Borsani L, Sergio F. N-Acetylcysteine as an antioxidant and disulphide breaking agent: the reasons why. Free Radic Res. 2018 Jul;52(7):751–762. doi: 10.1080/10715762.2018.1468564.

Aytan N, Choi JK, Carreras I, Brinkmann V, Kowall NW, Jenkins BG, Dedeoglu A. Fingolimod modulates multiple neuroinflammatory markers in a mouse model of Alzheimer’s disease. Sci Rep. 2016 Apr 27;6:24939. doi: 10.1038/srep24939.

Baslow MH. N-acetylaspartate in the vertebrate brain: metabolism and function. Neurochem Res. 2003 Jun;28(6):941–53. doi: 10.1023/a:1023250721185.

Belenguer P, Duarte JMN, Schuck PF, Ferreira GC. Mitochondria and the Brain: Bioenergetics and Beyond. Neurotox Res. 2019 Aug;36(2):219–238. doi: 10.1007/s12640-019-00061-7.

Brand A, Leibfritz D, Hamprecht B, Dringen R. Metabolism of cysteine in astroglial cells: synthesis of hypotaurine and taurine. J Neurochem. 1998 Aug;71(2):827–32. doi: 10.1046/j.1471-4159.1998.71020827.x.

Camargo RL, Batista TM, Ribeiro RA, Velloso LA, Boschero AC, Carneiro EM. Effects of taurine supplementation upon food intake and central insulin signaling in malnourished mice fed on a high-fat diet. Adv Exp Med Biol. 2013;776:93–103. doi: 10.1007/978-1-4614-6093-0_10.

Chiquita S, Ribeiro M, Castelhano J, Oliveira F, Sereno J, Batista M, Abrunhosa A, Rodrigues-Neves AC, Carecho R, Baptista F, Gomes C, Moreira PI, Ambrósio AF, Castelo-Branco M. A longitudinal multimodal in vivo molecular imaging study of the 3xTg-AD mouse model shows progressive early hippocampal and taurine loss. Hum Mol Genet. 2019 Jul 1;28(13):2174–2188. doi: 10.1093/hmg/ddz045.

Choy KH, Dean O, Berk M, Bush AI, van den Buuse M. Effects of N-acetyl-cysteine treatment on glutathione depletion and a short-term spatial memory deficit in 2-cyclohexene-1-one-treated rats. Eur J Pharmacol. 2010 Dec 15;649(1-3):224–8. doi: 10.1016/j.ejphar.2010.09.035.

de Paula GC, Brunetta HS, Engel DF, Gaspar JM, Velloso LA, Engblom D, de Oliveira J, de Bem AF. Hippocampal Function Is Impaired by a Short-Term High-Fat Diet in Mice: Increased Blood-Brain Barrier Permeability and Neuroinflammation as Triggering Events. Front Neurosci. 2021 Nov 4;15:734158. doi: 10.3389/fnins.2021.734158.

Deepmala, Slattery J, Kumar N, Delhey L, Berk M, Dean O, Spielholz C, Frye R. Clinical trials of N-acetylcysteine in psychiatry and neurology: A systematic review. Neurosci Biobehav Rev. 2015 Aug;55:294–321. doi: 10.1016/j.neubiorev.2015.04.015.

Duarte J.M.N., Cunha R.A., Carvalho R.A. (2007) Different metabolism of Glutamatergic and GABAergic compartments in superfused hippocampal slices characterized by nuclear magnetic resonance spectroscopy. Neurosci. 144, 1305–1313.

Duarte JMN, Carvalho RA, Cunha RA, Gruetter R. Caffeine consumption attenuates neurochemical modifications in the hippocampus of streptozotocin-induced diabetic rats. J Neurochem. 2009 Oct;111(2):368–79. doi: 10.1111/j.1471-4159.2009.06349.x.

Duarte JMN, Do KQ, Gruetter R. Longitudinal neurochemical modifications in the aging mouse brain measured in vivo by 1H magnetic resonance spectroscopy. Neurobiol Aging. 2014 Jul;35(7):1660–8. doi: 10.1016/j.neurobiolaging.2014.01.135. Epub 2014 Jan 31. PMID: 24560998.

Duarte JMN, Kulak A, Gholam-Razaee MM, Cuenod M, Gruetter R, Do KQ. N-acetylcysteine normalizes neurochemical changes in the glutathione-deficient schizophrenia mouse model during development. Biol Psychiatry. 2012 Jun 1;71(11):1006–14. doi: 10.1016/j.biopsych.2011.07.035. Epub 2011 Sep 25. PMID: 21945305.

Duarte JMN, Lei H, Mlynárik V, Gruetter R. The neurochemical profile quantified by in vivo 1H NMR spectroscopy. Neuroimage. 2012 Jun;61(2):342–62. doi: 10.1016/j.neuroimage.2011.12.038.

Duarte JMN, Skoug C, Silva HB, Carvalho RA, Gruetter R, Cunha RA. Impact of Caffeine Consumption on Type 2 Diabetes-Induced Spatial Memory Impairment and Neurochemical Alterations in the Hippocampus. Front Neurosci. 2019 Jan 9;12:1015. doi: 10.3389/fnins.2018.01015.

Duarte JMN, Xin L. Magnetic Resonance Spectroscopy in Schizophrenia: Evidence for Glutamatergic Dysfunction and Impaired Energy Metabolism. Neurochem Res. 2019 Jan;44(1):102–116. doi: 10.1007/s11064-018-2521-z.

Dwir D, Cabungcal JH, Xin L, Giangreco B, Parietti E, Cleusix M, Jenni R, Klauser P, Conus P, Cuénod M, Steullet P, Do KQ. Timely N-Acetyl-Cysteine and Environmental Enrichment Rescue Oxidative Stress-Induced Parvalbumin Interneuron Impairments via MMP9/RAGE Pathway: A Translational Approach for Early Intervention in Psychosis. Schizophr Bull. 2021 Oct 21;47(6):1782–1794. doi: 10.1093/schbul/sbab066.

El Idrissi A. Taurine improves learning and retention in aged mice. Neurosci Lett. 2008 May 2;436(1):19–22. doi: 10.1016/j.neulet.2008.02.070.

Falach-Malik A, Rozenfeld H, Chetboun M, Rozenberg K, Elyasiyan U, Sampson SR, Rosenzweig T. N-Acetyl-L-Cysteine inhibits the development of glucose intolerance and hepatic steatosis in diabetes-prone mice. Am J Transl Res. 2016 Sep 15;8(9):3744–3756.

Figueroa AL, Figueiredo H, Rebuffat SA, Vieira E, Gomis R. Taurine Treatment Modulates Circadian Rhythms in Mice Fed A High Fat Diet. Sci Rep. 2016 Nov 18;6:36801. doi: 10.1038/srep36801.

Garcia-Serrano AM, Duarte JMN. Brain Metabolism Alterations in Type 2 Diabetes: What Did We Learn From Diet-Induced Diabetes Models? Front Neurosci. 2020 Mar 20;14:229. doi: 10.3389/fnins.2020.00229.

Garcia-Serrano AM, Mohr AA, Philippe J, Skoug C, Spegel P, Duarte JMN (2022) Cognitive impairment and metabolite profile alterations in the hippocampus and cortex of male and female mice exposed to a fat and sugar-rich diet are normalized by diet reversal. Aging Dis 10.14336/AD.2021.0720 (in press)

Gbd 2019 Dementia Forecasting Collaborators. Estimation of the global prevalence of dementia in 2019 and forecasted prevalence in 2050: an analysis for the Global Burden of Disease Study 2019. Lancet Public Health. 2022 Jan 6:S2468-2667(21)00249–8. doi: 10.1016/S2468-2667(21)00249-8. Epub ahead of print. PMID: 34998485.

Guerreiro RJ, Gustafson DR, Hardy J. The genetic architecture of Alzheimer’s disease: beyond APP, PSENs and APOE. Neurobiol Aging. 2012 Mar;33(3):437–56. doi: 10.1016/j.neurobiolaging.2010.03.025. Epub 2010 Jul 1. PMID: 20594621; PMCID: PMC2980860.

Haider S, Sajid I, Batool Z, Madiha S, Sadir S, Kamil N, Liaquat L, Ahmad S, Tabassum S, Khaliq S. Supplementation of Taurine Insulates Against Oxidative Stress, Confers Neuroprotection and Attenuates Memory Impairment in Noise Stress Exposed Male Wistar Rats. Neurochem Res. 2020 Nov;45(11):2762–2774. doi: 10.1007/s11064-020-03127-7.

Harris JL, Choi IY, Brooks WM. Probing astrocyte metabolism in vivo: proton magnetic resonance spectroscopy in the injured and aging brain. Front Aging Neurosci. 2015 Oct 28;7:202. doi: 10.3389/fnagi.2015.00202.

Iorio E, Podo F, Leach MO, Koutcher J, Blankenberg FG, Norfray JF. A novel roadmap connecting the 1H-MRS total choline resonance to all hallmarks of cancer following targeted therapy. Eur Radiol Exp. 2021 Jan 15;5(1):5. doi: 10.1186/s41747-020-00192-z.

Kim HY, Kim HV, Yoon JH, Kang BR, Cho SM, Lee S, Kim JY, Kim JW, Cho Y, Woo J, Kim Y. Taurine in drinking water recovers learning and memory in the adult APP/PS1 mouse model of Alzheimer’s disease. Sci Rep. 2014 Dec 12;4:7467. doi: 10.1038/srep07467.

Kim KS, Jang MJ, Fang S, Yoon SG, Kim IY, Seong JK, Yang HI, Hahm DH. Anti-obesity effect of taurine through inhibition of adipogenesis in white fat tissue but not in brown fat tissue in a high-fat diet-induced obese mouse model. Amino Acids. 2019 Feb;51(2):245–254. doi: 10.1007/s00726-018-2659-7.

Livingston G, Huntley J, Sommerlad A, Ames D, Ballard C, Banerjee S, Brayne C, Burns A, Cohen-Mansfield J, Cooper C, Costafreda SG, Dias A, Fox N, Gitlin LN, Howard R, Kales HC, Kivimäki M, Larson EB, Ogunniyi A, Orgeta V, Ritchie K, Rockwood K, Sampson EL, Samus Q, Schneider LS, Selbæk G, Teri L, Mukadam N. Dementia prevention, intervention, and care: 2020 report of the Lancet Commission. Lancet. 2020 Aug 8;396(10248):413–446. doi: 10.1016/S0140-6736(20)30367-6.

Lizarbe B, Soares AF, Larsson S, Duarte JMN (2019). Neurochemical Modifications in the Hippocampus, Cortex and Hypothalamus of Mice Exposed to Long-Term High-Fat Diet. Front Neurosci. 12:985. doi: 10.3389/fnins.2018.00985.

Louzada PR, Paula Lima AC, Mendonca-Silva DL, Noël F, De Mello FG, Ferreira ST. Taurine prevents the neurotoxicity of beta-amyloid and glutamate receptor agonists: activation of GABA receptors and possible implications for Alzheimer’s disease and other neurological disorders. FASEB J. 2004 Mar;18(3):511–8. doi: 10.1096/fj.03-0739com.

Lynch J. W. (2004). Molecular structure and function of the glycine receptor chloride channel. Physiological reviews, 84(4), 1051–1095. https://doi.org/10.1152/physrev.00042.2003

McLean FH, Campbell FM, Sergi D, Grant C, Morris AC, Hay EA, MacKenzie A, Mayer CD, Langston RF, Williams LM. Early and reversible changes to the hippocampal proteome in mice on a high-fat diet. Nutr Metab (Lond). 2019 Aug 23;16:57. doi: 10.1186/s12986-019-0387-y.

More J, Galusso N, Veloso P, Montecinos L, Finkelstein JP, Sanchez G, Bull R, Valdés JL, Hidalgo C, Paula-Lima A. N-Acetylcysteine Prevents the Spatial Memory Deficits and the Redox-Dependent RyR2 Decrease Displayed by an Alzheimer’s Disease Rat Model. Front Aging Neurosci. 2018 Dec 6;10:399. doi: 10.3389/fnagi.2018.00399.

Qizilbash N, Gregson J, Johnson ME, Pearce N, Douglas I, Wing K, Evans SJW, Pocock SJ. BMI and risk of dementia in two million people over two decades: a retrospective cohort study. Lancet Diabetes Endocrinol. 2015 Jun;3(6):431–436. doi: 10.1016/S2213-8587(15)00033-9.

Ribeiro RA, Santos-Silva JC, Vettorazzi JF, Cotrim BB, Mobiolli DD, Boschero AC, Carneiro EM. Taurine supplementation prevents morpho-physiological alterations in high-fat diet mice pancreatic β-cells. Amino Acids. 2012 Oct;43(4):1791–801. doi: 10.1007/s00726-012-1263-5.

Shahidi S, Zargooshnia S, Asl SS, Komaki A, Sarihi A. Influence of N-acetyl cysteine on beta-amyloid-induced Alzheimer’s disease in a rat model: A behavioral and electrophysiological study. Brain Res Bull. 2017 May;131:142–149. doi: 10.1016/j.brainresbull.2017.04.001.

Shen FC, Weng SW, Tsao CF, Lin HY, Chang CS, Lin CY, Lian WS, Chuang JH, Lin TK, Liou CW, Wang PW. Early intervention of N-acetylcysteine better improves insulin resistance in diet-induced obesity mice. Free Radic Res. 2018 Dec;52(11-12):1296–1310. doi: 10.1080/10715762.2018.1447670.

Shivaraj MC, Marcy G, Low G, Ryu JR, Zhao X, Rosales FJ, Goh EL. Taurine induces proliferation of neural stem cells and synapse development in the developing mouse brain. PLoS One. 2012;7(8):e42935. doi: 10.1371/journal.pone.0042935.

Sonnay S, Gruetter R, Duarte JMN. How Energy Metabolism Supports Cerebral Function: Insights from 13C Magnetic Resonance Studies In vivo. Front Neurosci. 2017 May 26;11:288. doi: 10.3389/fnins.2017.00288.

Takado Y, Takuwa H, Takuya U, Takahashi M, Ono M, Maeda J, Shimojo M, Nitta N, Shibata S, Aoki I, Sahara N, Suhara T, Higuchi M (2018) Correlations between brain metabolites and tau protein accumulation assessed by 1H-MRS and tau PET in Alzheimer’s disease model mice. Proc Intl Soc Mag Reson Med 26, 4976. http://indexsmart.mirasmart.com/ISMRM2018/PDFfiles/4976.html

Tondo M, Wasek B, Escola-Gil JC, de Gonzalo-Calvo D, Harmon C, Arning E, Bottiglieri T. Altered Brain Metabolome Is Associated with Memory Impairment in the rTg4510 Mouse Model of Tauopathy. Metabolites. 2020 Feb 14;10(2):69. doi: 10.3390/metabo10020069.

Vitvitsky V, Garg SK, Banerjee R. Taurine biosynthesis by neurons and astrocytes. J Biol Chem. 2011 Sep 16;286(37):32002–10. doi: 10.1074/jbc.M111.253344.

Zhang H, Huang M, Gao L, Lei H. Region-specific cerebral metabolic alterations in streptozotocin-induced type 1 diabetic rats: an in vivo proton magnetic resonance spectroscopy study. J Cereb Blood Flow Metab. 2015 Nov;35(11):1738–45. doi: 10.1038/jcbfm.2015.111.

Zhou Y, Holmseth S, Guo C, Hassel B, Höfner G, Huitfeldt HS, Wanner KT, Danbolt NC. Deletion of the γ-aminobutyric acid transporter 2 (GAT2 and SLC6A13) gene in mice leads to changes in liver and brain taurine contents. J Biol Chem. 2012 Oct 12;287(42):35733–35746. doi: 10.1074/jbc.M112.368175.

